# A likelihood method for estimating present-day human contamination in ancient DNA samples using low-depth haploid chromosome data

**DOI:** 10.1101/594481

**Authors:** J. Víctor Moreno-Mayar, Thorfinn Sand Korneliussen, Anders Albrechtsen, Jyoti Dalal, Gabriel Renaud, Rasmus Nielsen, Anna-Sapfo Malaspinas

## Abstract

1

**Motivation:** The presence of present-day human contaminating DNA fragments is one of the challenges defining ancient DNA (aDNA) research. This is especially relevant to the ancient *human* DNA field where it is difficult to distinguish endogenous molecules from human contaminants due to their genetic similarity. Recently, with the advent of high-throughput sequencing and new aDNA protocols, hundreds of ancient human genomes have become available. Contamination in those genomes has been measured with computational methods often developed specifically for these empirical studies. Consequently, some of these methods have not been implemented and tested while few are aimed at low-depth data, a common feature in aDNA datasets.

**Results:** We develop a new X-chromosome-based maximum likelihood method for estimating present-day human contamination in low-depth sequencing data. We implement our method for general use, assess its performance under conditions typical of ancient human DNA research, and compare it to previous nuclear data-based methods through extensive simulations. For low-depth data, we show that existing methods can produce unusable estimates or substantially underestimate contamination. In contrast, our method provides accurate estimates for a depth of coverage as low as 0.5× on the X-chromosome when contamination is below 25%. Moreover, our method still yields meaningful estimates in very challenging situations, *i.e.*, when the contaminant and the target come from closely related populations or with increased error rates. With a running time below five minutes, our method is applicable to large scale aDNA genomic studies.

**Availability and implementation:** The method is implemented in **C++ and R** and is freely available in https://github.com/sapfo/contaminationX.

## 2 Introduction

Having plagued the field since its inception (Zischler et al., 1995), contamination is one of the defining features of ancient DNA (aDNA). While DNA extracted from present-day specimens is mostly endogenous, aDNA extracts are a mixture of low levels of damaged and fragmented endogenous DNA often dwarfed by higher amounts of contaminant DNA (Orlando et al., 2015). In recent years, high-throughput sequencing technologies have substantually contributed to advancing the field by randomly retrieving DNA fragments present in the extract, *i.e.*, including the shorter, damaged endogenous ones. Nevertheless, the problem of contamination has persisted, and affects all laboratories (Wall and Kim, 2007; Champlot et al., 2010; Llamas et al., 2017; Der Sarkissian et al., 2015; Pääbo et al., 2004; Willerslev and Cooper, 2005; Sampietro et al., 2006; Gilbert et al., 2005).

Contaminant DNA is expected to have either an environmental (*e.g.* soil microbes) or a human origin *e.g.* people involved in excavation, extraction or sample handling (Sampietro et al., 2006; Llamas et al., 2017). As aDNA sequencing data is routinely mapped to a reference genome that is closely related to the study organism (Schubert et al., 2012), identifying environmental contamination by means of sequence identity is relatively straightforward. However, for human samples, human contamination can be particularly pernicious as endogenous and exogenous DNA molecules are highly similar. Moreover, this type of contamination is problematic as it could lead to spurious evolutionary inferences (Wall and Kim, 2007; Racimo et al., 2016). Consequently, a number of methods for quantifying contamination in aDNA data have emerged during the last decade. Existing methods rely on either haploid chromosomes (*e.g.*, the mitochondrial DNA (mtDNA) (Fu et al., 2013; Green et al., 2008; Renaud et al., 2015) and the X-chromosome in males (Rasmussen et al., 2011)) or diploid autosomes (Racimo et al., 2016).

### MtDNA-based methods

Mitochondrial DNA is often present in multiple almost identical copies in a given cell and is considerably shorter than the nuclear genome. As such, mtDNA has been historically easier to target and sequence compared to the nuclear genome (Higuchi et al., 1984; Krings et al., 1997). Hence, the first computational methods to measure contamination were tailored to this short molecule for which a high depth of coverage is often achieved. In general, methods based on haploid genomic segments (*e.g.*, mtDNA) rely on the expectation that there is a single DNA sequence type per cell. Thus, multiple alleles at a given site would be the result of either contamination, *post-mortem* damage, sequencing or mapping error.

Currently, there are three common mitochondrial DNA-based methods that require a high coverage mtDNA consensus sequence. Green et al. (Green et al., 2008), estimated mtDNA contamination in a Neanderthal sample by counting the number of reads that did not support the mtDNA consensus (assumed to be the endogenous sequence) at sites where the consensus differed from a worldwide panel of mtDNAs (‘fixed derived sites’). Later, Fu et al. (Fu et al., 2013) introduced a method focused on modelling the observed reads as a mixture of the mtDNAs in a panel containing the endogenous sequence while co-estimating an error parameter. Importantly, these methods did not take into account the complexity of inferring the endogenous ‘consensus’ mtDNA sequence. Thus, a subsequent method (Schmutzi) sought to jointly infer the endogenous mitogenome while estimating present-day human contamination via the incorporation of the intrinsic characteristics of endogenous aDNA fragments into the model (Renaud et al., 2015).

### Autosomes-based methods

Sequencing high depth ancient nuclear genomes remains challenging. Therefore, mtDNA-based contamination estimates have been used as a proxy for overall contamination (Allentoft et al., 2015). Yet, different mitochondrial-to-nuclear DNA ratios in the endogenous source and the human contaminant(s) may lead to inaccurate conclusions (Furtwängler et al., 2018). While the source of this difference has yet to be identified, accurate methods based on nuclear data are needed to estimate the level of human contamination which may have an impact on downstream analyses (Renaud et al., 2016). Indeed, most studies rely on nuclear data to answer key biological questions. A recent method (DICE) aims at estimating present-day human contamination for nuclear data (Racimo et al., 2016). It does so by co-estimating contamination, sequencing error, and demography based on autosomal data. This method generally requires an intermediate depth of coverage (at least 3×) and produces more accurate results when the sample and the contaminant are genetically distant (*e.g.* different species or highly differentiated populations).

### X-chromosome-based methods and a novel approach

In 2011, Rasmussen et al. (Rasmussen et al., 2011) estimated the contamination level in whole genome sequencing data from a male Aboriginal Australian based on the X-chromosome using a maximum likelihood method. Similar to mtDNA-based methods, this method relies on the fact that the X-chromosome is hemizygous in males. The mathematical details of the method used in that study were described in the supplementary information. However, while this method could in principle also perform well for low depth data, its performance was not assessed in detail.

In this work, we propose a new maximum likelihood method (implemented in **C++** and R) relying on ‘relatively long’ haploid chromosomes potentially sequenced at low depth of coverage (such as the X-chromosome in male humans). We present the mathematical details of our method, perform extensive simulations and analyze real data to compare it to existing nuclear-based methods. To do so, we also implement the method by (Rasmussen et al., 2011) (see Sections 3.3 and 6 for a discussion on the fundamental differences between methods). We measure the performance of the methods by quantifying the accuracy of the contamination estimates and assess the effect of a) varying levels of contamination, b) varying depth of coverage, c) the ancestry of the endogenous and the contaminant populations and d) additional error in the endogenous data. We show that our method performs particularly well for low-depth data compared to other methods. It can accurately estimate present-day human contamination for male samples that are likely to be candidates for further evolutionary analysis (*i.e.* when contamination is <25%) when the X-chromosome depth of coverage is as low as 0.5×. Moreover, our implementation is fast and scalable.

## 3 Methods

We assume we have collected high-throughput whole genome sequence (WGS) data from a sample that contains DNA from two different sources; DNA belonging to one individual of interest (the ‘endogenous’ DNA or ‘endogenous individual’), and DNA from contaminating individuals. We want to estimate the fraction *c* of DNA that belongs to the contaminant individuals versus the individual of interest. We assume that the individual of interest and the contaminants belong to the same species but they can belong to different populations. We denote the contaminating population by *Pop*_*c*_. Given the high-throughput nature of the data, each site along the genome can be covered by multiple sequencing reads or alleles. The data has been mapped to a reference genome which includes a haploid chromosome (*e.g.*, the X-chromosome for human males). Across all chromosomes, a fraction *c* of the reads belong to the contaminants while the rest (1 − *c*) belong to the endogenous individual.

For haploid chromosome(s), we expect that the individual of interest will carry only one allele at each site, and we rely on this idea to estimate *c*, the contamination fraction. As discussed above, observing multiple alleles at a given site can be due to either sequencing error, *post-mortem* DNA degradation, mapping errors or contamination.

### 3.1 Assumptions and notation

We rely on the availability of population genetic data (allele frequencies) from a ‘reference panel’ from a number of populations including *Pop*_*c*_. We assume that (1) the panel includes data at *L* polymorphic sites; (2) there are four possible bases (*A, C, G* and *T*) at every site but only two are naturally segregating across populations (we have bi-allelic sites) (3) we know the population allele frequencies of *Pop*_*c*_ perfectly; (4) the endogenous individual carries either naturally segregating alleles with equal probability (see discussion); (5) there are no mapping errors, hence multiple alleles will only be due to error (sequencing or *post-mortem* damage) or contamination; (6) all observed sequencing reads are independent draws from a large pool of DNA sequences.

At every site *i*, we denote 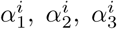 and 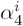 the potential alleles that we can observe, with 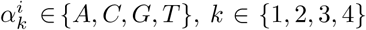 and *i* ∈ {1, …, *L*}. To simplify the presentation, we will assume that at all sites 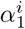 and 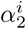 occur naturally in the population (bi-allelic sites), while 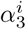 and 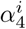 can be observed because of sequencing error or damage. For each site included in the reference panel, there is a single true allele carried by the individual of interest (the endogenous allele), where there could be also contaminant alleles. We call these the ‘endogenous allele’ 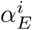 and the ‘contaminant allele(s)’ 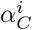. The frequencies of the segregating alleles across sites in the contaminating population (*Pop*_*c*_) will be denoted by the matrix 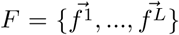, where 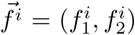 are the frequencies of the alleles 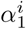 and 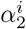 in that population at site *i*.

We further assume that errors affect all bases equally and that they occur independently across reads and across bases within a read. The probability of having an error from base *a* ∈ {A, C, G, T} to base *b* ∈ {A, C, G, T} is given by the matrix Γ = {*γ*_*ab*_}. While this can be easily generalized, in our current implementation, we will set *γ*_*ab*_ = *ϵ/*3 if *a* ≠ *b* and therefore *γ*_*aa*_ = (1 − ϵ) *∀ a, b* ∈ {A, C, G, T}. In other words we assume that all types of mutations are equally likely. See Section 3.4 for details on the estimation of Γ.

Finally, we summarise the data with the total counts of 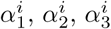 and 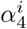 alleles at every site and we label those counts 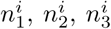 and 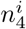 with 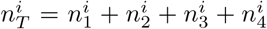. We extend this notation to also keep track of multiple alleles, so for instance 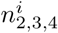 is the number of 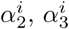 or 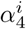 alleles.

### 3.2 Model description – a likelihood approach

Let us now assume that 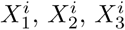 and 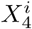 are random variables keeping track of the number of 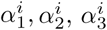 and 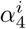 alleles that can be observed in the data at site *i*. We also write 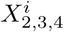, for instance, for the number of 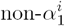 alleles. We can then denote 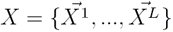 the random variable summarizing the high-throughput observed data across polymorphic sites, with 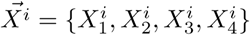.

We will first compute the probability of the counts of a given allele at site *i* given the allele frequencies *F* in the contaminating population, the contamination rate *c* and the error matrix Γ, which we then use for computing the likelihood of the full data (see below, Equation 41). We start by conditioning on the endogenous allele. We have that:

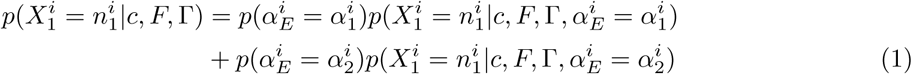

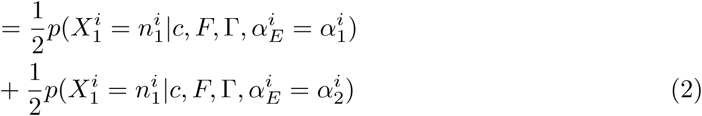

since there is a single true endogenous allele at each site and we have assumed that the endogenous individual a priori carries either allele with equal probability. If the pool of sequencing reads we draw from is large enough, which is likely to be the case with high-throughput data, we have that each draw is identically distributed for a given endogenous allele. Hence, given an endogenous allele, the alleles we draw at each site follow a binomial distribution. Relabeling:

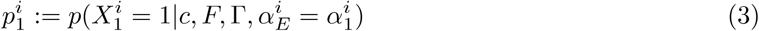

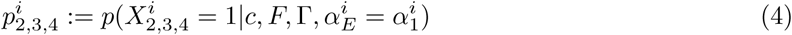

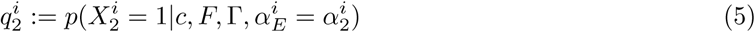

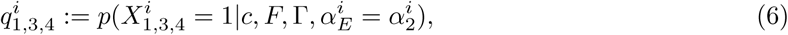

The probability of seeing 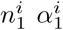 alleles in the data assuming the endogenous allele is 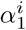 and that we have a total of 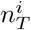 sequenced reads at that site is given by:

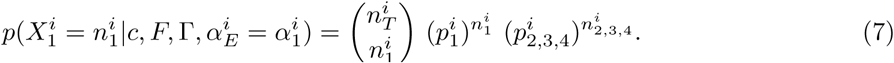

Similarly, if the endogenous is 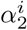, we have that:

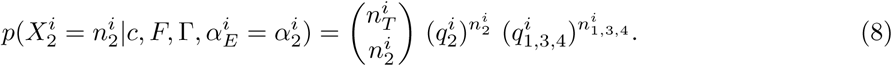

We can now compute the probability of 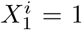, that is the probability of observing one 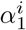 allele in the sequencing data. We will momentarily drop the index *i* to simplify the presentation. Let us first assume that the true endogenous allele is *α*_1_ (*i.e.*, we first compute *p*_1_). By conditioning on the source of the observed allele being either the endogenous (‘endo’) or a contaminant (‘cont’) individual, we have that:

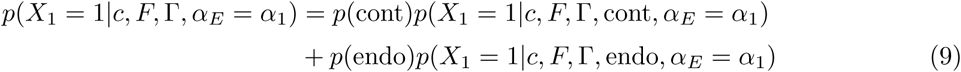

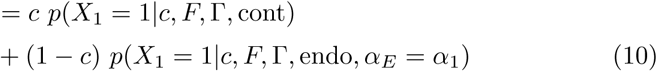

In the contaminant case, we then condition on either of the naturally segregating alleles:

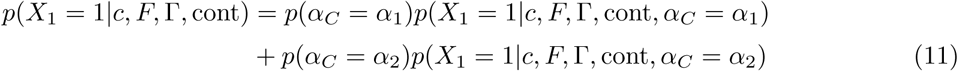

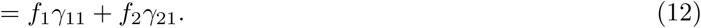

While for an endogenous draw we have:

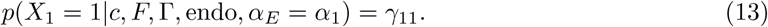

By substituting the equations above into equation (10) we have that:

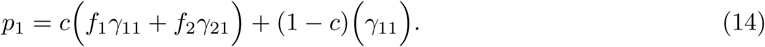

There are indeed two ways to draw an *α*_1_ allele. First, we could draw a read from a contaminating individual. This individual belongs to population *Pop*_*c*_ and there is therefore a probability *f*_1_ that it carries that allele, and *f*_2_ that it carries the alternative allele *α*_2_. If it carries *α*_1_, we would need no error to occur (*γ*_11_). While if the contaminant carries *α*_2_, it would need to mutate to *α*_1_ (*γ*_21_). Second, we could draw a read from the endogenous individual. Since we have assumed that the endogenous individual carries an *α*_1_ allele, it should remain *α*_1_, *i.e.*, no error (*γ*_11_). We can similarly obtain all other three equations for the probability of observing an *α*_2_, *α*_3_ or *α*_4_ allele:

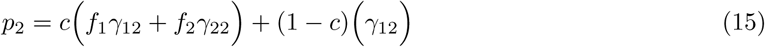

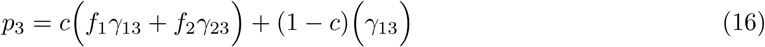

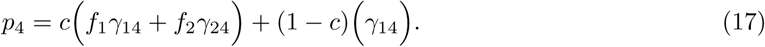

The equivalent expression for observing non-*α*_1_ alleles is simply

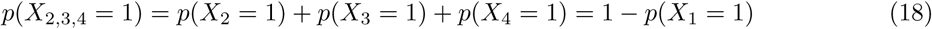

since it is not possible to draw simultaneously two alleles. We then have that:

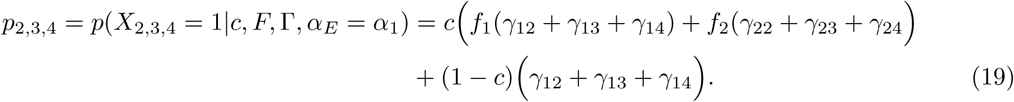

Conditioning on the endogenous allele being *α*_2_ and following a similar logic, we have for the *q*_*k*_ equations:

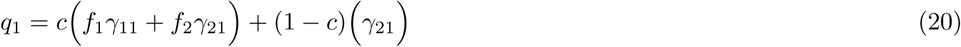

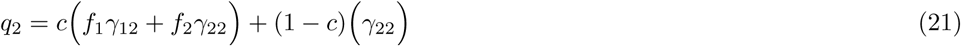

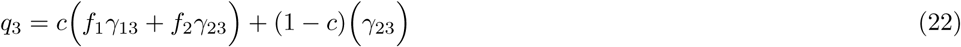

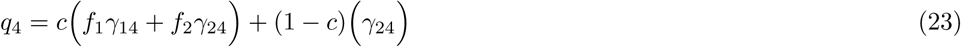

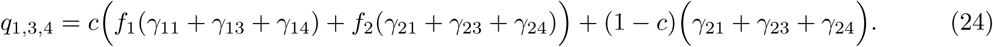

The first part of the *q*_*k*_ equations, corresponding to the contaminant read case, is identical to the first part of the *p*_*k*_ equations 14, 15, 16, and 17. For the second part, which corresponds to the endogenous read case, we can simply invert indices 1 and 2 to recover the second part of the *p*_*k*_ equations. We can simplify all equations further since in our implementation we have *γ*_*aa*_ = (1 − *ϵ*) and *γ*_*ab*_ = *ϵ/*3 *∀ a, b* ∈ {A, C, G, T} with *a ≠ b*. Adding now the *i* index, we have for the 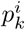:

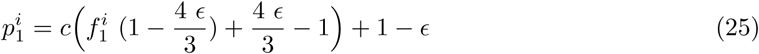

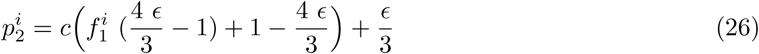

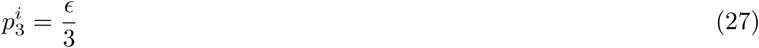

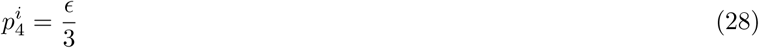

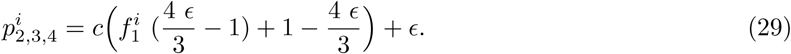

Note that we can further simplify those expressions by using 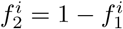:

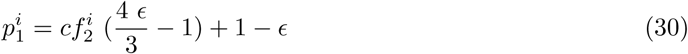

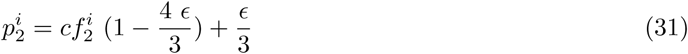

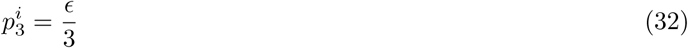

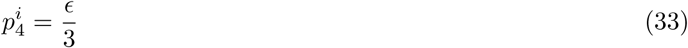

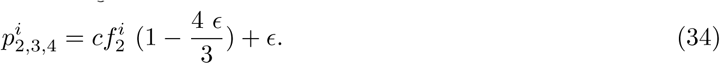

And for the 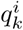:

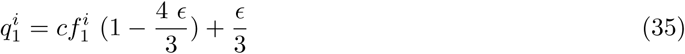

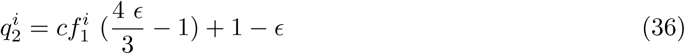

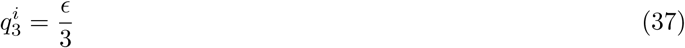

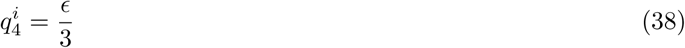

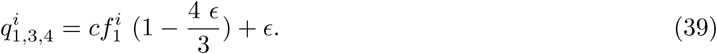

#### Likelihood function – ‘Two-consensus’

We will filter the data so that a read only covers one polymorphic site. In other words, since the reads are assumed to be independent from each other, each site is also independent. Assuming the error rates are known (see below), the likelihood function for the parameter *c* can be written as:

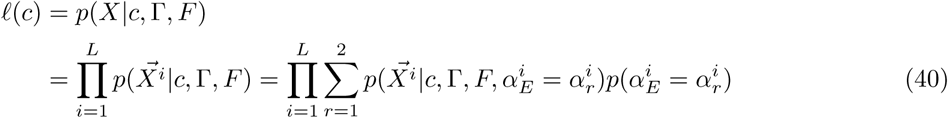

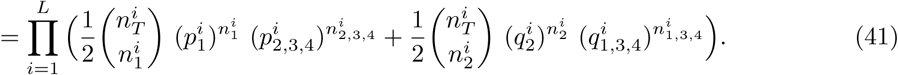

We can then find the value *c* (*ĉ*_*mle*_) that maximizes *ℓ*(*c*) (*i.e.* the maximum likelihood estimate, mle).

### 3.3 Previous related approach – ‘One-consensus’

The method we propose above is related to one that was described in the supplementary material of (Rasmussen et al., 2011). The key difference, beside the consideration that a contaminant allele may also have errors, is that Rasmussen et al. assumed that at each polymorphic site, the most prevalent allele in the sequencing data was the true endogenous allele. Without loss of generality, we can call this allele *α*_1_. In other words, we assume that at every site *p*(*α*_*E*_ = *α*_1_) = 1 and *p*(*α*_*E*_ = *α*_2_) = 0. Denoting 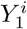 the number of consensus *α*_1_ alleles and 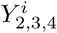 the number of non-consensus alleles, we have that:

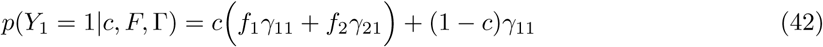

Similarly, for *Y*_2,3,4_, we have that:

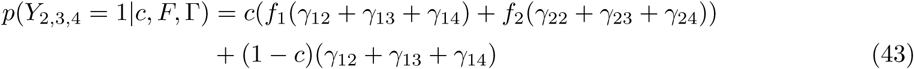

Finally, denoting 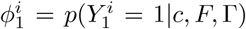 and 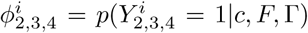, and expressing the errors rates in terms of *ϵ*, we have as above:

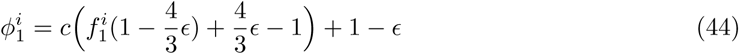

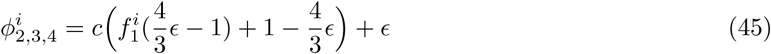

While the likelihood function becomes:

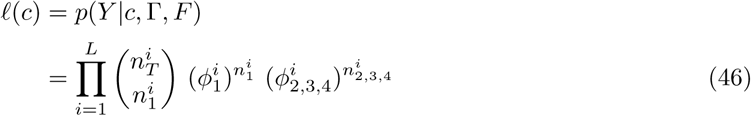

since *p*(*α*_*E*_ = *α*_2_) = 0. We call this approach the ‘One-consensus’ method since the ‘consensus’ allele is assumed to be the truth; accordingly, we will call our new approach the ‘Two-consensus’ method since we integrate over both segregating alleles and assume that either can be the true endogenous (consensus) allele at a particular site.

### 3.4 Estimating error rates

To infer the contamination rate *c*, we first obtain a point estimate of *E* by considering the flanking regions of the polymorphic sites following (Rasmussen et al., 2011). Specifically, we assume that the sites neighboring a polymorphic site *i* in the reference panel are fixed across all populations – including population *Pop*_*c*_ and are given by the most prevalent allele at each of those sites. Without loss of generality we can assume *α*_1_ = *α*_*C*_ = *α*_*E*_ for all flanking sites. We label the flanking sites *i*_*j*_ where, *e.g., i*_−2_ is the second site to the left of site *i* (*i*_0_ is site *i*). We assume that non-*α*_1_ alleles at those neighboring sites are solely due to error. In other words when *j ≠* 0, we have that 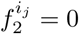, and hence 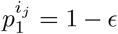 and 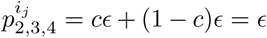 (Equations (30) and (34)). We consider the counts of non-*α*_1_ alleles at *s* sites left and right of the polymorphic sites. Having assumed that (i) reads are independent of each other, (ii) bases within a read are independent from each other, we have:

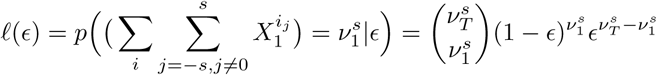

where 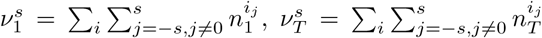. To infer the contamination rate, we then substitute the error rate in Equation 41 by the maximum likelihood estimate of the error rate obtained at the flanking regions across polymorphic sites, which is simply: 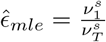. Note that by default we set *s* = 4, *i.e.*, we consider four sites left and right of the polymorphic site to compute the error rate.

### 3.5 Standard error

To compute the standard error for the inferred parameter, we consider a block jackknife approach that we apply to the likelihood approach. Specifically we split the haploid chromosome into *M* blocks, each corresponding to one of the *L* sites (we have *M ≤ L*). For each *m* = 1*…M* we leave one block *m* out and compute 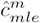 over the remaining data. We estimate the standard error for the estimate using the following relationship:

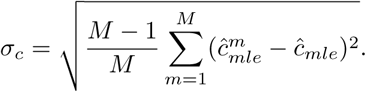

Under some regularity conditions, the 95% confidence interval for our contamination rate is then *ĉ±* 2*s*_*c*_.

### 3.6 Implementation

Our method is implemented as two separate steps. First, the counts of bases are tabulated for a sample provided by the user as a bam file of mapped reads. This is done within the software ANGSD (Korneliussen et al., 2014) which allows to filter the data efficiently and is implemented in c++. The contamination estimates are obtained in the second step based on the output from step one along with a file containing information about the reference population (polymorphism data from a reference panel). This step is implemented in R. The documentation along with a description and explanation of options and output are found on the following website: https://github.com/sapfo/contaminationX. The human reference population allele frequency panels used in this study are available there as well.

## 4 Performance assessment

To evaluate our method’s performance in practice, we carried out simulations with parameters typical of human aDNA experiments. Although we focused on humans, the method is expected to be equally applicable to other species for which polymorphism data are available. In particular, we assessed the effect on the estimates of 1. the contamination fraction, 2. the depth of coverage, 3. the genetic distance between the sample and the contaminant, 4. the genetic distance between the contaminant and the reference panel assumed to be the contaminating population, and 5. the error rate. In addition, we compared our method to two existing methods based on nuclear data; namely, our implementation of the ‘One-consensus’ method by Rasmussen et al. (2011) and DICE by Racimo et al. (2016). In all cases, we simulated sequencing data by sampling and ‘mixing’ mapped reads from publicly available genomes in known proportions while controlling for the depth of coverage (DoC).

### 4.1 General simulation framework and settings

For all experiments described below we used our method with the following settings: -d 3, -e 20 (*i.e.*, filtering for sites with a minimum DoC of 3 and a maximum of 20) and maxsites=1000 (resampling at most 1,000 blocks for the block jackknife procedure). To compare methods and parameter values, we computed the root mean square error (*RMSE*), the bias and the range for a set of *k* contamination estimates from simulated data *Ĉ* = {*ĉ*_1_, *ĉ*_2_, *…, ĉ*_*k*_} and an expected contamination fraction *c*_*exp*_ (where applicable) as follows:

1. 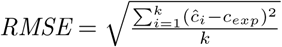

2. 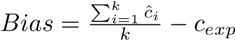

3. *Range* = *max*(*Ĉ*) *-min*(*Ĉ*)

For all experiments where we estimated *RMSE, Bias* and *Range*, we simulated 100 replicates for each parameter combination.

### 4.2 Test genomes and reference panels

We considered Illumina whole genome sequencing data from a subset of the present-day individuals reported in (Meyer et al., 2012). We included data from six male individuals ranging in DoC between 19.9× and 26.7×: a Yoruba (HGDP00927), a Karitiana (HGDP00998), a Han (HGDP00778), a Papuan (HGDP00542), a Sardinian (HGDP00665), and a French (HGDP00521). All data were pre-processed, mapped and filtered following (Malaspinas et al., 2014).

We considered ten populations from the HapMap project as potential proxies for *Pop*_*c*_. Those populations represent broad scale worldwide variation (Altshuler et al., 2010). We filtered each panel by removing: 1) all sites located in the pseudoautosomal region of the human X chromosome (parameters -b 5000000 -c 154900000 discard the first 5Mb and last *∼*370Kb of the human X chromosome, following Ensembl GRCh37 release 95); 2) all sites with a minor allele frequency lower than 0.05 (-m 0.05); 3) all variable sites located less than 10 bp away from another variable site. The number of remaining sites after filtering each panel is shown in Table 1.

**Table 1:** Reference allele frequency panels used for estimating contamination. *Number of single nucleotide polymorphism (SNPs) included for each population after applying the filtering described in the text. Data were downloaded from http://hapmap.ncbi.nlm.nih.gov/downloads/frequencies/2010-08_phaseII+III/allele_freqs_chrX_CEU_r28_nr.b36_fwd.txt.gz

| Population | Number of sites | Number of sites (filtered)* | Number of individuals |
| --- | --- | --- | --- |
| HapMap_ASW | 38,703 | 31,324 | 90 |
| HapMap_CEU | 73,562 | 58,190 | 180 |
| HapMap_CHB | 67,307 | 51,494 | 90 |
| HapMap_GIH | 34,158 | 26,098 | 100 |
| HapMap_JPT | 64,290 | 49,715 | 91 |
| HapMap_LWK | 39,992 | 31,119 | 100 |
| HapMap_MEX | 34,360 | 23,190 | 90 |
| HapMap_MKK | 37,935 | 29,612 | 180 |
| HapMap_TSI | 33,928 | 25,097 | 100 |
| HapMap_YRI | 89,604 | 72,546 | 180 |

### 4.3 One- vs Two-consensus methods and reasonable parameter range for c

We first explored the contamination fractions for which our method yields informative estimates. To do so, we sampled 1× data from a Yoruba individual and ‘contaminated’ these with data from a French individual at increasing contamination rates {0.01, 0.05, 0.1, …, 0.45, 0.50}. Note that by design, our method cannot distinguish between ‘symmetric’ contamination fractions, *e.g.*, 0.2 from 0.8. For this exploratory analysis, we simulated five replicates for each contamination rate and used the HapMap CEU reference panel as a proxy for the allele frequencies in the contaminant population. For each simulation, we estimated the contamination fraction using the ‘One-consensus’ (Rasmussen et al., 2011) and the ‘Two-consensus’ methods.

The results are shown on Figure 1a. We observed that the estimated contamination rates matched the simulated rates qualitatively for both methods as long as the contamination fraction was below 0.25 (see below for a discussion relative to the bias). In addition, the ‘Two-consensus’ method provided more accurate results especially when contamination was high. Given both methods failed at estimating very large contamination fractions accurately, we simulate data with contamination rates between 0.01 and 0.25 for subsequent analyses.

**Figure 1:**
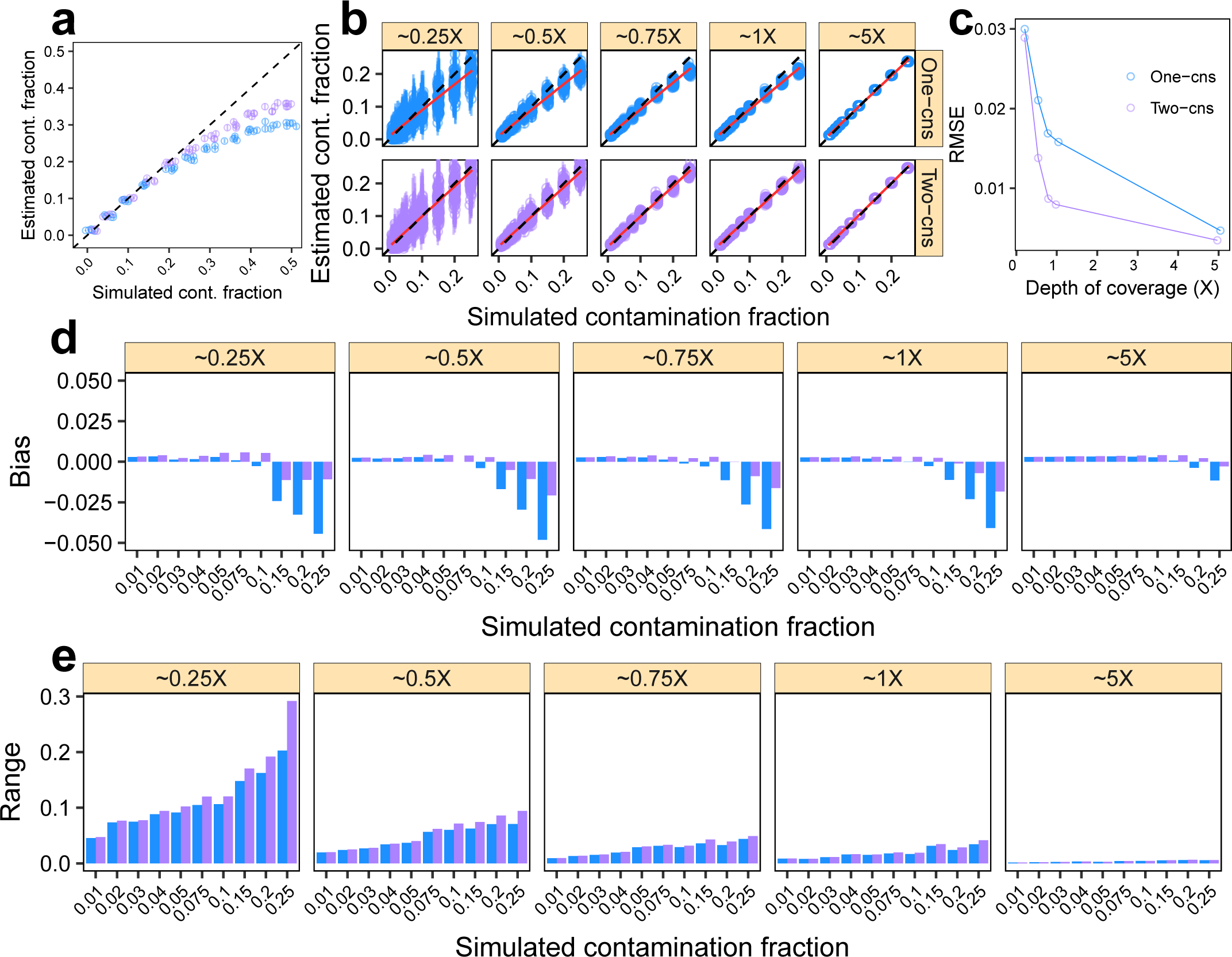
Parameter range for *c* and effect of the DoC for the One- and Two-consensus methods. We simulated data as described in Sections 4.3 and 4.4 to explore the contamination fractions and DoC for which our method yields informative estimates: we ‘contaminated’ a Yoruba with a French individual with increasing contamination fractions while controlling for the DoC. a,b. contamination estimates for each replicate (points) and corresponding 95% confidence intervals (vertical bars). The dashed lines indicate the expected values and the red lines a linear regression. c. *RMSE* for each DoC, combining the results across simulated contamination fractions in b. d. *Bias* for each DoC and contamination fraction combination. e. *Range* for each DoC and contamination fraction combination. Results for the ‘One-consensus’ and ‘Two-consensus’ methods are shown in blue and purple, respectively across all panels.

### 4.4 One- vs Two-consensus methods and depth of coverage

We carried out a similar simulation experiment to determine the broad effect of the DoC on the estimates of the ‘One-consensus’ and the ‘Two-consensus’ methods. In this case, we sampled sequencing data at varying DoC {0.25×, 0.5×, 0.75×, 1×, 5×} with increasing contamination rates {0.01, 0.02, 0.03, 0.04, 0.05, 0.075, 0.1, 0.2, 0.25}. Results are summarized in Figure 1b,c,d,e.

We found that both methods yielded estimates close to the truth, especially when the contamination fraction was within the simulation range [0.01, 0.25] and the DoC was *≥*0.5× (Figure 1b). As expected, the range of the estimates increased with lower DoC and higher contamination fractions (Figure 1e). The *RMSE* also decreased with higher DoC, while we observed that this decrease slowed down between 0.75× and 1×.

We observed that both methods slightly overestimated contamination for true contamination fractions *<*0.1 and underestimated it for values *>*0.1. Importantly, the downward bias for large contamination fractions and the RMSE (specially between 0.5× and 5×) were substantially lower for the ‘Two-consensus’ method compared to the ‘One-consensus’ one. This difference in bias is intuitive and follows from the mathematical details of each of the methods (see also discussion). Thus, since the ‘Two-consensus’ approach performed equally well for higher DoC and outperformed the previous method with lower DoC, we see no advantage in using the ‘One-consensus’ method and focus hereafter on characterizing the ‘Two-consensus’.

### 4.5 Comparison with DICE

We compared the performance of our method to DICE, an autosomal data-based method for co-estimating contamination, sequencing error, and demography (Racimo et al., 2016). We carried out simulations as detailed above and we ‘contaminated’ an ancient Native American genome (Anzick1) (Rasmussen et al., 2014) with data from a present-day French individual. In this case, we used an ancient individual to favor DICE, which jointly estimates the error rate and contamination fraction. We ran DICE with the two-population model using the 1000 Genomes Project Phase III CEU allele frequencies as a proxy for the frequencies of the putative contaminant and the YRI frequencies to represent the ‘anchor’ population. We let the MCMC algorithm run for 100,000 steps and discarded as burn-in the first 10,000 steps. We used the coda R package to obtain 95% posterior credibility intervals. For our method we used the parameters detailed in Section 4.1. We summarise the results for this comparison in Figure 2.

**Figure 2:**
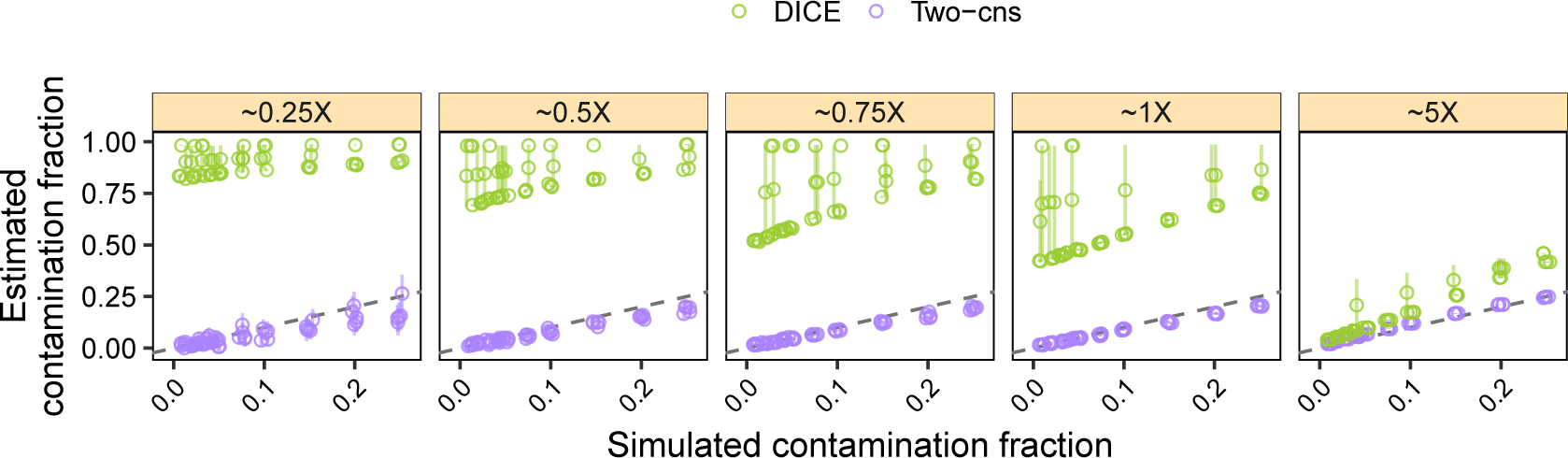
Simulation results comparing our method to DICE. We simulated data as described in Section 4.5 and estimated contamination across five replicates using our method (purple) and DICE (green). We ‘contaminated’ the Anzick1 ancient Native American genome with a French individual at increasing contamination fractions while controlling for the DoC. Vertical bars correspond to 95% confidence intervals for the Two-consensus method and to 95% credible intervals for DICE. The dashed line indicates the expected values. Note that the simulated DoC corresponds to the autosomal DoC for DICE and the X-chromosome DoC for our method.

In agreement with the simulations based on present-day data in the previous section, we observed that our method yielded accurate estimates for a DoC as low as 0.5× and for true contamination fractions below 0.25. In contrast, in most cases, we observed that DICE did not converge to a value close to the simulated contamination fraction for a DoC *≤* 1 but instead vastly overestimated contamination. Whereas DICE started to yield useful estimates at 5×, our method provided more accurate estimates than DICE for all simulated cases. These results suggest that for low depth data (*≤* 5×) the ‘Two-consensus’ method should be used to estimate contamination.

### 4.6 Lowest bound on depth of coverage for the Two-consensus method

To get a sense of the minimal amount of data necessary to obtain accurate estimates with our method, we carried out simulations for a more fine-grained range of DoC {0.1×, 0.2×, 0.3×, 0.4×, 0.5×, 0.6×, 0.7×, 0.8×, 0.9× and 1×}. Results are summarised in Figure 3. In agreement with results presented in Section 4.4, we observed that across simulations, the estimates closely matched the truth from 0.2× onward (see linear regression). Similarly, the *RMSE* sharply decreased at 0.2× while it qualitatively saturated from 0.5× onward. In other words, our estimates are already meaningful for a DoC as low as 0.2×, and become quite accurate for a DoC *≥*0.5×. Based on these results, when the reference panel used for estimation is a close representative of the contaminant population (see also Section 4.8), we recommend to use our method to determine if a sample or library is highly contaminated (contamination *>*25%), or to estimate the contamination fraction when contamination is between 0 and 25%.

**Figure 3:**
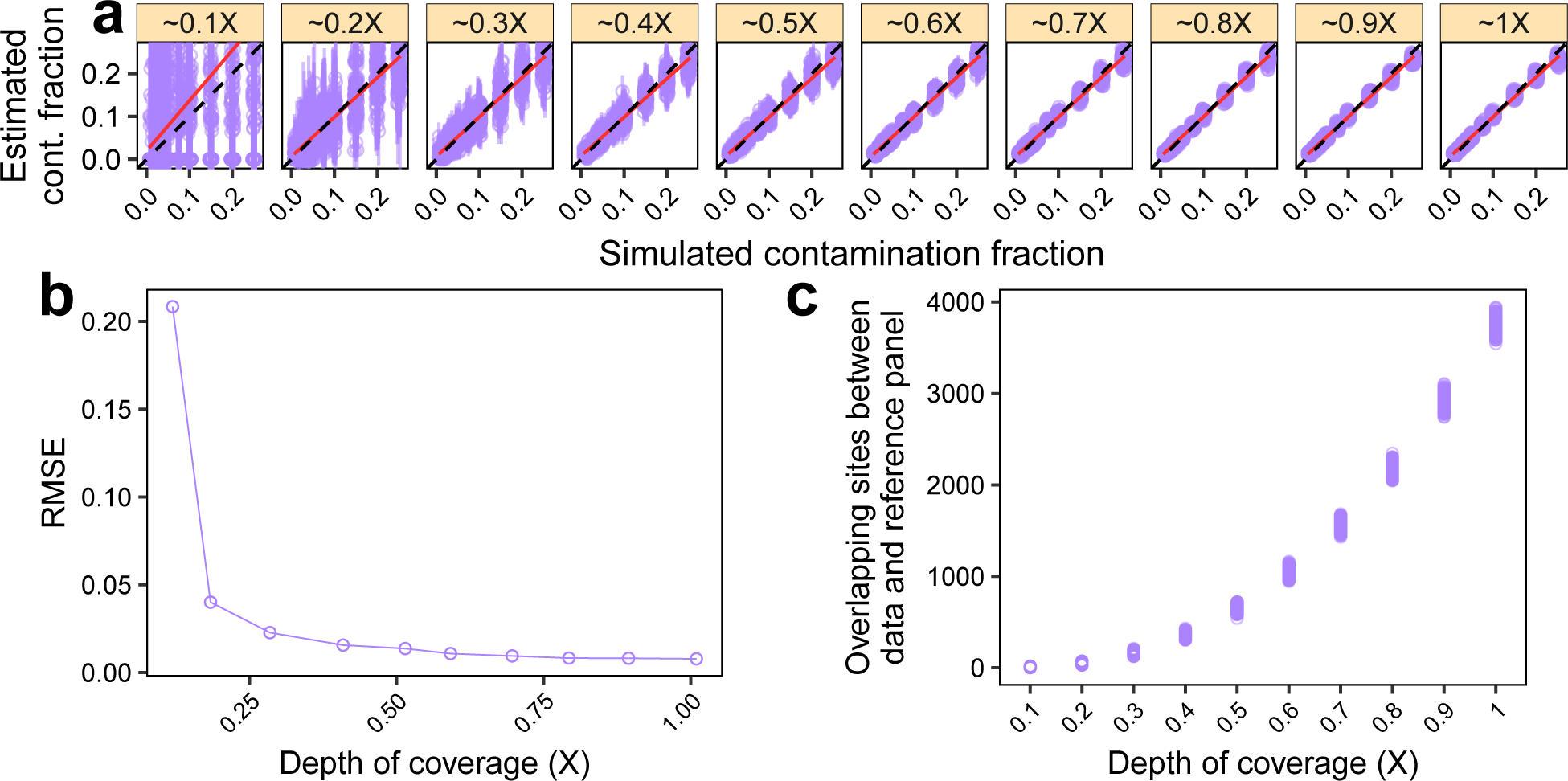
Minimum required depth of coverage (DoC). We simulated data as described in Section 4.4, but we considered an additional range of low DoC {0.01×, 0.02×, …, 1×}. a. contamination estimates for each replicate (points) and corresponding 95% confidence intervals (vertical bars). Dashed lines indicate the expected values and red lines show a linear regression. b: *RMSE* for each DoC, combining the results across contamination fractions from a. c. Number of overlapping sites between the simulated data and the contaminant population panel (HapMap CEU in this case) after applying the filters detailed in Section 4.1.

### 4.7 The effect of the genetic distance between the endogenous and the contaminant individuals

While we do not consider the ancestry of the endogenous individual in our model, intuitively, estimating the contamination fraction should be easier when the endogenous and contaminant individuals are more distantly related. To get further insights into this intuition, we sampled sequencing data from five individuals (a Yoruba, a Karitiana, a Han, a Papuan and a Sardinian) and contaminated them with data from a French individual. We used the same depth of coverage and contamination fraction settings described in Section 4.4 and used the HapMap CEU reference panel to estimate the contamination fraction. We explored the relationship of the contamination estimates and the ‘allele sharing distance’ between the X-chromosome consensus sequences from the five individuals and the French contaminant. We defined the allele sharing distance as the number of differences between the French and each individual’s consensus, divided by the number of non-missing sites for each pair.

Results are shown in Figure 4. We obtained a very similar picture across simulated endogenous individuals. Indeed, the *RMSE*, the bias and the range of the estimates vary as a function of the DoC with qualitativly little effect from the genetic distance between the contaminant and the endogenous individual. As such, our method seemingly performs equally well regardless of the ancestry of the endogenous individual, even for cases where contaminant and endogenous are closely related (*e.g.* a Sardinian individual contaminated with a French individual).

**Figure 4:**
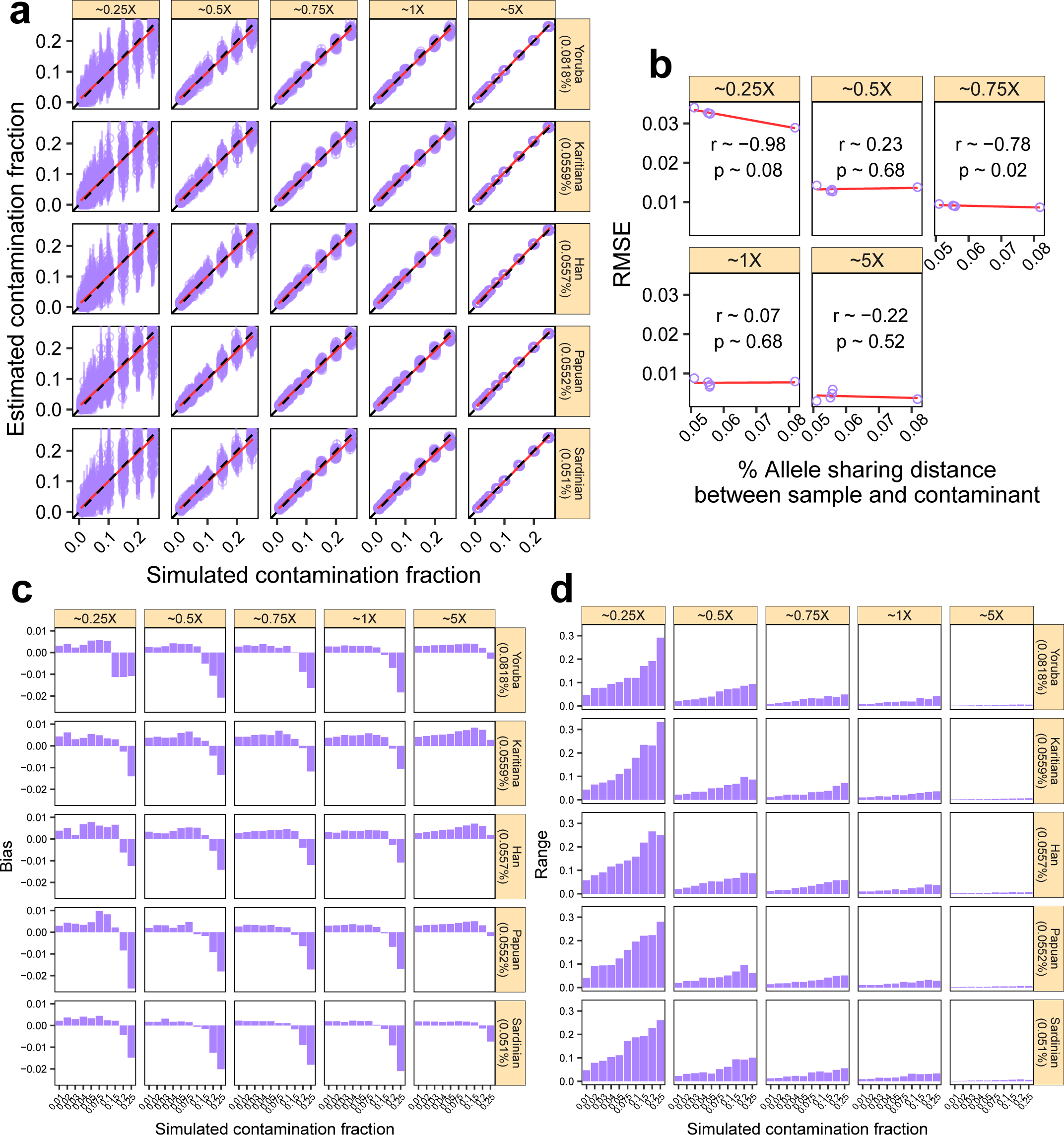
The effect of the genetic distance between the endogenous and the contaminant individuals. We considered five individuals (Yoruba, Karitiana, Han, Papuan, Sardinian) and ‘contaminated’ them with a French individual (Section 4.7). We simulated data with increasing contamination fractions while controlling for the DoC. a. contamination estimates for each replicate (points) and corresponding 95% confidence intervals (vertical bars). Dashed lines indicate the expected values and red lines show a linear regression. The allele sharing distance between each sample and the contaminant is indicated in parentheses. b. *RMSE* for each DoC as a function of the allele sharing distance between the five samples and the contaminant, combining the results across contamination fractions in a. We show the Pearson correlation coefficient for each DoC. c. *Bias* for each DoC, sample and contamination fraction combination. d. *Range* for each DoC, sample and contamination fraction combination.

### 4.8 The effect of the genetic distance between the simulated contaminant and the reference panel used for inferring contamination

For this experiment, we sampled data from a Sardinian individual and contaminated it with data from a French individual. We applied the same depth of coverage and contamination fraction settings from the above experiments and used ten different reference populations from the HapMap project as proxies for *Pop*_*c*_: ASW, CEU, CHB, GIH, JPT, LWK, MEX, MKK, TSI and YRI, to estimate the contamination fraction. To get an indicative value for the distance between the reference HapMap panel and the contaminant, we estimated the genetic distance between the X-chromosome consensus sequence from the contaminant French individual and each reference population. We defined this distance as 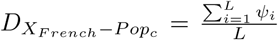 where *L* is the total number of sites included in the reference population *Pop*_*c*_ (assumed to be the contaminant) and *ψ*_*i*_ is the frequency of the allele carried by the contaminant individual *X* (French in this case), at locus *i*. Note that we only considered the sites that are included in all reference panels to compute this distance. Results are shown in Figure 5.

**Figure 5:**
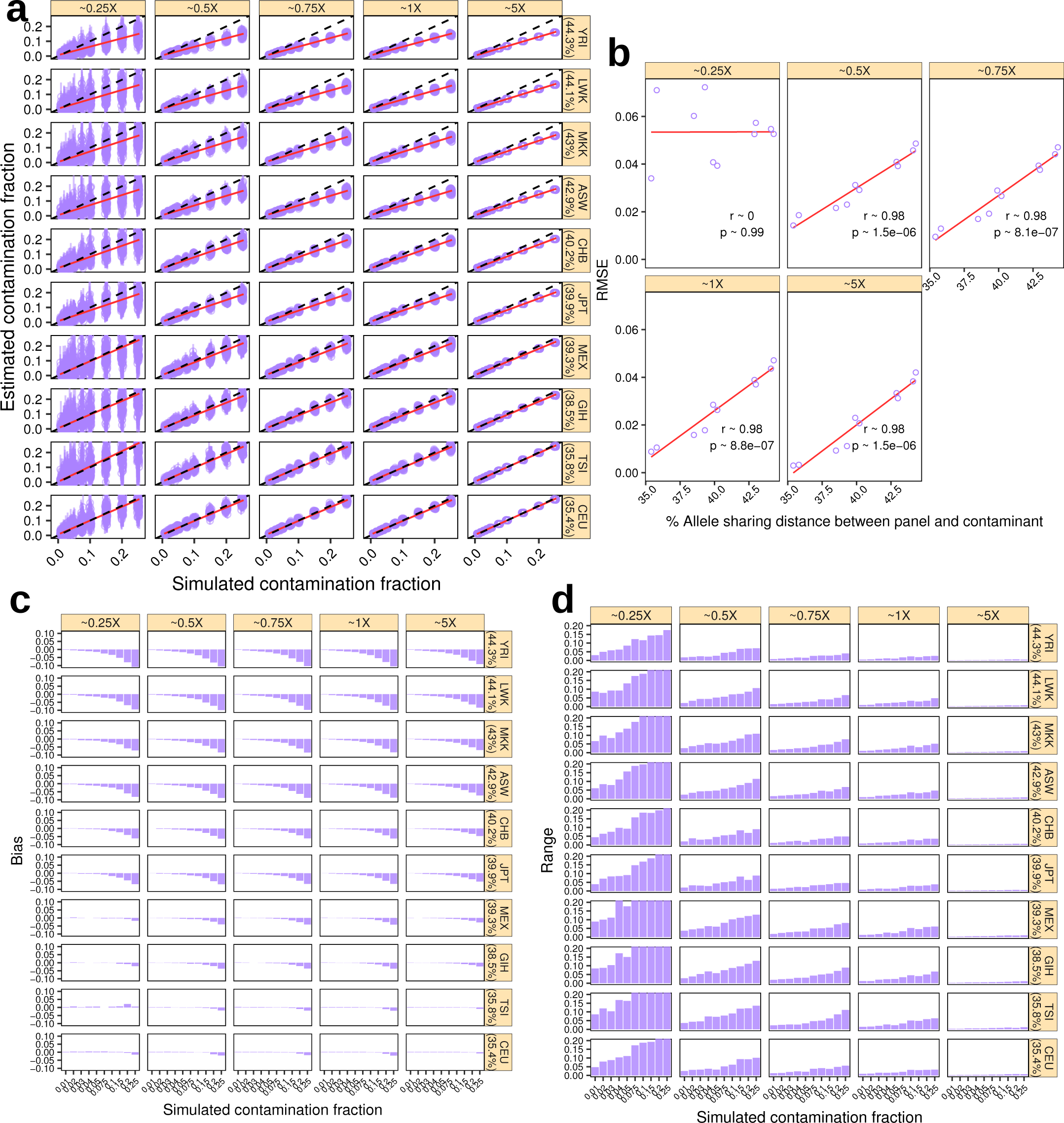
The effect of the distance between the reference population (*Pop*_*c*_) and the contaminant. We simulated data as described in Section 4.7. We considered the ten reference populations described in Table 1 and ‘contaminated’ a Sardininan with a French individual. We simulated data with increasing contamination fractions while controlling for the DoC. a. contamination estimates for each replicate (points) and corresponding 95% confidence intervals (vertical bars). Dashed lines indicate the expected values and red lines show a linear regression. The genetic distance between the reference panel 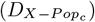 is indicated in parentheses. b. *RMSE* for each DoC as a function of 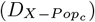, combining the results across contamination fractions in a. We show the Pearson correlation coefficient for each DoC. c. *Bias* for each DoC, sample and contamination fraction combination. d: *Range* for each DoC, sample and contamination fraction combination.

We found that misspecifying the contaminant population led to an underestimation of the contamination fraction (Figure 5a). In fact, as indicated by the strong correlation between the *RMSE* and the genetic distance 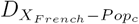, worse ‘guesses’ of the contaminant ancestry resulted in worse estimates. This correlation was similar across all tested DoC but 0.25×. We observed a downward bias for larger simulated contamination fractions that increased with 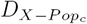. Although the overall effect could be deemed relatively small (*e.g., RMSE <*0.05 with the HapMap YRI panel), if the contaminating population is not known, we recommend comparing results obtained through different reference populations. Note that one could also use this observation to make a qualitative statement about the ancestry of the contaminant individual assuming several reference populations are available (see, for example, (Rasmussen et al., 2015)).

### 4.9 The effect of differential error rates in the endogenous and contaminant individuals

We assessed the effect of varying the error rates in the endogenous sequencing data by simulating data as detailed above. However, in this case, we added errors to the Yoruba reads at a constant rate ϵ ∈ {0.005, 0.01, 0.02, 0.05, 0.1} by using a transition matrix Γ = *γ*_*ab*_ analogous to the one used for error rate estimation. Results are summarized in Figure 6. Qualitatively, although there is a significant positive correlation between the *RMSE* and the error (Figure 6b), the overall effect is small, except for the extreme cases of 5% and 10% added error, where we observe a systematic overestimation of contamination. Yet, we note that current second generation sequencing platforms such as the Illumina HiSeq, have substantially lower error rates, *e.g.*, sequencing error rates in the modern human genome dataset from (Meyer et al., 2012) have been estimated to be between 0.03 and 0.05% (Malaspinas et al., 2014). The apparent innocuousness of additional small amounts of error, is likely due to the fact that error affects all sites (variable and neighboring) uniformly in our model, but also that the error rate is smaller than the explored range of contamination rate (except for 5% and 10% added error).

**Figure 6:**
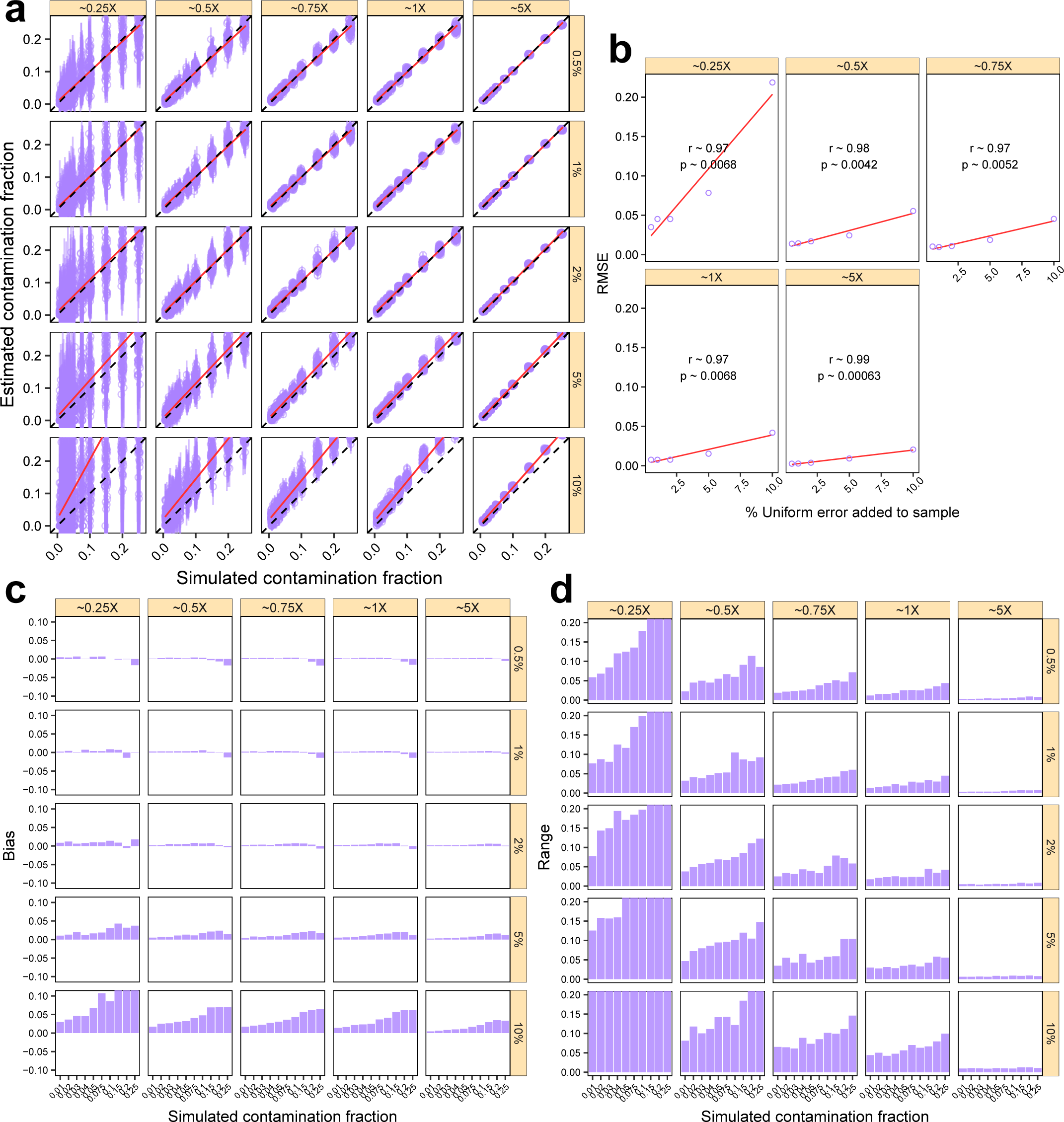
The effect of differential error rates in the endogenous individual. We simulated data as described in Section 4.4 and added error increasingly to the Yoruba individual. a. contamination estimates for each replicate (points) and corresponding 95% confidence intervals (vertical bars). Dashed lines indicate the expected values and red lines show a linear regression. Added error rates are indicated to the right of each panel. b. *RMSE* for each DoC as a function of the added error. We show the Pearson correlation coefficient for each DoC. c. *Bias* for each DoC, added error and contamination fraction combination. d. *Range* for each DoC, added error and contamination fraction combination.

We note that the observed error structure for aDNA is different from our simulations. In particular the error is not independent of the position across reads. For example, C to T and G to A misincorporations tend to accumulate towards the reads’ termini (Briggs et al., 2007). However, we expect damage-derived error to be uniform across polymorphic sites, in the sense that segregating and neighboring sites are equally likely to be damaged. Therefore, we do not expect aDNA damage to inflate contamination estimates differently from how uniform error does. We note, however, that if variable sites are more error-prone than neighboring sites due to sequence-intrinsic features, contamination may be overestimated. In Section 4.5, we showed that contamination estimates for simulations involving real aDNA data are qualitatively similar to those obtained for simulations with present-day data.

## 5 Running time

We explored the running time of our method implementation using a machine with 24 2.8 GHz Intel Xeon cores. The data parsing step for 5× X-chromosome datasets was always below 3 minutes. Following data parsing, the raw contamination estimate is obtained nearly instantaneously. Thus, the step that requires the largest amount of time is the calculation of the standard error. Since we use a jackknife approach this will have a running time of 𝒪^2^ in the number of sites. Therefore, the actual running time will depend on the depth of coverage and the number of polymorphic sites in the reference panel. Using the parameters detailed in Section 4.1, we estimated the contamination fraction in the *∼*14× Anzick1 genome (Rasmussen et al., 2014) with a joint running time of approximately three minutes for the parsing and estimation steps.

## 6 Discussion

We present here a new method for efficiently estimating contamination in low depth high-throughput sequencing data based on information from haploid chromosomes. To assess whether our method can be used in challenging situations typical of aDNA research, we tested it through realistic simulations and assess its performance. Note that our simulations involved a single contaminating individual —a realistic assumption in our view. Yet, our method can in principle handle multiple contaminants from *Pop*_*c*_, which we anticipate would improve our method’s performance as the simulations would match the implemented model more closely. Our simulations suggest that our method can correctly flag highly contaminated samples from male individuals that are unlikely to be useful in evolutionary analyses (*c ≥*25%), and outputs an accurate contamination estimate for male samples with lower amounts of contamination (*c <*25%).

Based on the results above, we show that provided one can approximatively guess the contaminant reference population, our estimates will be meaningful even when DoC is as low as 0.2× and essentially unbiased when contamination is below 15%. We also show that our method is easily scalable since the running time is below five minutes for a depth of coverage as high as 10X (on the X-chromosome). Based on these features, we regard our method as an adequate and practical tool for screening large numbers of aDNA male samples and related libraries to get a sense of candidates for follow-up analyses. Indeed, aDNA studies have transitioned to the genomic era with single studies sometimes including whole genomes (Damgaard et al., 2018) or genome-wide SNP data (Olalde et al., 2018) from hundreds of individuals. However, most ancient samples carry low proportions of endogenous DNA and the resulting depth of coverage for a given shotgun experiment is often quite low for laboratories working with a finite budget. Thus, prioritizing resources on promising samples is often a key aspect of human aDNA research.

We have shown that typical sequencing error rates and the genetic distance between the endogenous and contaminant individuals do not affect the accuracy of our estimates. However, we found that misspecifying the contaminant population leads to underestimation (*Bias <*0.1). In particular, while the method is still able to detect contamination, this issue is more pronounced when contamination is *>*10%. In practice, our method flags contaminated samples with estimates *>*10% and we recommend that the user takes a conservative approach: explore several potential contaminant populations and report the highest estimate. Note that a high error rate could in principle impact the accuracy, but our simulations suggest this would lead to an overestimation of contamination, *i.e.*, our method would be conservative in this case.

Finally, we show that our method outperforms the previously published nuclear genome data-based methods ‘One-consensus’ (Rasmussen et al., 2011) and DICE (Racimo et al., 2016). It outperforms them in particular for low depth data (*<*5×) and when contamination is above 10%. The main difference between the One- and Two-consensus is that for the latter we do not assume that the true endogenous allele is the observed consensus at each site. This assumption is particularly wrong for low depth data, even when filtering for sites with at least 3 reads. Since we show the ‘Two-consensus’ method is more accurate across the parameter space we explored, our new method is a better choice. In contrast, DICE offers additional functionality by co-estimating contamination, error rates and demography using autosomal data. Thus, while DICE is not useful for screening (or estimating contamination for) low depth samples, an appropriate protocol would comprise an initial screening using the ‘Two-consensus’ method, followed by further deeper sequencing. If the resulting DoC is *>*5× DICE could be used to co-estimate contamination and the demography.

## Acknowledgements

We thank Philip L. F. Johnson, Fernando Racimo, Frédéric Michaud, Florian Clemente, M. Thomas P. Gilbert and Ludovic Orlando for helpful discussion.

